# A new species of subgenus *Typhlodromus* (*Anthoseius*) De Leon (Acari: Phytoseiidae), and phytoseiid mites intercepted in samples imported from 12 different countries to Taiwan

**DOI:** 10.1101/2020.09.20.305656

**Authors:** Jhih-Rong Liao, Chyi-Chen Ho, Chiun-Cheng Ko

**Affiliations:** Department of Entomology, National Taiwan University, Taipei City 10617, Taiwan; Taiwan Acari Research Laboratory, Taichung City, Taiwan

**Keywords:** Phytoseiidae, plant quarantine, new species, checklist.

## Abstract

Global trade has increased the invasion risk of exotic organisms and damaged agricultural and natural ecosystems. The Bureau of Animal and Plant Health Inspection and Quarantine (BAPHIQ) handles quarantine services of animal- and plant-associated pests and diseases in Taiwan. The predatory mite family Phytoseiidae (Acari: Mesostigmata) is a well-known group due to the potential use of certain species as biocontrol agents for small phytophagous pests. Some species are available in commercial markets and frequently used in biological control in many agricultural systems especially in greenhouse crops. However, exotic biological control agents may interfere with native or naturalized populations of predatory mites and they may threaten indigenous populations via intraguild predation. The present study aims to provide the phytoseiid mite species found in plant quarantine from 2006–2013. Twenty-five species belonging two subfamilies and eight genera were found in samples imported to Tawan from twelve countries, including one new species *Typhlodromus* (*Anthoseius*) *ueckermanni* **sp. nov.** from South Africa. The checklist provides distribution, remarks, and also an identification key for all species.

## Introduction

International trade has allowed exotic species to move across geographical barriers and invade to new areas. Exotic organisms may have negative effects on agricultural and natural ecosystems (Hulme 2009). Numerous exotic natural enemies have been introduced and provide successful pest control. However, few studies have focused on the negative effects of introduced biological control agents (van Lenteren *et al*. 2003). The family Phytoseiidae (Acari: Mesostigmata) is a famous group in terms of their potential as biological control agents, and contains more than 2,700 species (including synonyms) worldwide (Chant & McMurtry 2007; Demite *et al*. 2020). Certain phytoseiids are available commercial markets to be used in biological control of spider mites, eriophyid mites, and small soft bodied insects such as thrips and whiteflies, and thus provide a valuable contribution to the agroecosystem (McMurtry *et al*. 2013). However, exotic biological control agents may interfere with native or naturalized populations of predatory mites and they may threaten indigenous populations via intraguild predation. For example, *Harmonia axyridis* (Pallas) is a famous generalist predator for hemipteran pests, however, now it is considered as an invasive species in many countries where it has been spreading rapidly and declined the populations of native species since the early years of the 21st century (Brown *et al*. 2018). In addition, Sato & Mochizuki (2011) reported that two exotic phytoseiids (*Neoseiulus cucumeris* and *Amblyseius swirskii*) were introduced to Japan, have advantage over an indigenous species, *Gynaeseius liturivorus* (Ehara) via intrguild predation. In this regard, the determination of predatory mites in plant quarantine samples imported from foreign countries is crucial for the evaluation of their environmental risk over natural population.

This study provides a checklist and an identification key of 25 phytoseiid species found during plant quarantine into Taiwan from 2006–2013. The checklist contains 121 records of phytoseiid mites from 12 countries-Australia, Canada, Chile, France, Israel, Japan, Malaysia, the Netherlands, New Zealand, Peru, Thailand, and the United States and 34 plant species. The study provides species distribution, specimen information, and remarks. Additionally, one new species *Typhlodromus* (*Anthoseius*) *ueckermanni* **sp. nov.,** was described and illustrated based on the specimens found in the plant material imported from South Africa is reported.

## Material and Methods

Mite specimens examined in this study were collected in plant quarantine by inspectors of Bureau of Animal and Plant Health Inspection and Quarantine during 2006–2013. Specimens were mounted in Hoyer’s medium, examined under an optical microscope (Olympus BX51). Measurements were taken using a stage-calibrated ocular micrometers and ImageJ 1.47 (Schneider *et al*. 2012). All measurements were provided in micrometers (μm) and holotype measurements are shown in bold type for new species, followed by their mean and range (in parenthesis). The dorsal shield lengths were measured from anterior to posterior margins along the midline and the widths measured at *j6* and *S4* levels. The sternal shield lengths and widths were taken from anterior to posterior margins along the midline and at broadest level, respectively. The genital shield widths were taken from broadest level. The ventrianal shield lengths were taken from anterior to posterior margins along the midline including cribrum and the shield widths measured at *ZV2* and anus levels. The general terminology used for morphological descriptions in this study follows that of Chant & McMurtry (2007). The notation for idiosomal setae follows that of Lindquist & Evans (1965) and Lindquist (1994), as adapted by Rowell *et al*. (1978) and Chant & Yoshida-Shaul (1992). The notation for solenostomes and lyrifissures is based on Athias-Henriot (1975). Specimens were deposited in TARL (Taiwan Acari Research Laboratory, Taichung City, Taiwan).

## Results

### Identification key to phytoseiid mites in plant quarantine in Taiwan based on female specimens

1. Dorsal setae *z3* and *s6* absent … Amblyseiinae … 2

Dorsal setae *z3* and *s6* present … Typhlodrominae … 16

2. Setae *S2* and *S4* absent … *Phytoseiulus persimilis*

Setae *S2* and *S4* present … 3

3. Ratio of setae *s4*:*Z1* > 3.0:1.0 … 4

Ratio of setae *s4*:*Z1* < 3.0:1.0 … 10

4. Seta *J2* absent … genus *Proprioseiopsis* … 5

Seta *J2* present … genus *Amblyseius* … 6

5. Spermatheca calyx cup-shaped … *Proprioseiopsis asetus*

Spermatheca calyx saccular … *Proprioseiopsis neotropicus*

6. Seta *z4* at least as long as 2/3 distance between its base and that of seta *s4* … *Amblyseius sinautus* Seta *z4* short or minute not as long as 2/3 distance between its base and that of seta *s4* … 7

7. Female ventrianal shield vase-shaped … *Amblyseius largoensis*

Female ventrianal shield usually pentagonal … 8

8. Calyx of spermatheca tubular … *Amblyseius tamatavensis*

Calyx of spermatheca dish-, cup, bell- or V-shaped … 9

9. Movable digit with two teeth … *Amblyseius tsugawai*

Movable digit with three teeth … *Amblyseius andersoni*

10. Spermatheca with atrium deeply forked … 11

Spermatheca with atrium not forked … 13

11. Atrium of spermatheca broader than based of calyx … *Neoseiulus makuwa*

Atrium of spermatheca as wide as calyx … 12

12. Length of *St* IV relative longer, ca. 58–74 µm … *Neoseiulus barkeri*

Length of *St* IV relative shorter, ca. 34 µm … *Neoseiulus loxus*

13. Spermatheca with a neck between calyx and atrium … *Neoseiulus bicaudus*

Spermtaheca without neck between calyx and atrium … 14

14. Movable digit of chelicerae with one tooth … *Neoseiulus cucumeris*

Movable digit of chelicerae with three teeth … 15

15. Macrosetae present only on tarsus IV … *Neoseiulus californicus*

Macrosetae present on genu, tibia and tarsus IV … *Neoseiulus cucumeris*

16. Seta *z3* absent … *Galendromus occidentalis*

Seta *z3* present … 17

17. Setae *S4* and *JV4* absent … 18

Setae *S4* and *JV4* present … genus *Typhlodromus* … 19

18. Setae z4 inserted mesad of setae *z2* and *S4* … *Meyerius immutatus*

Setae *z4* inserted in line with setae *z2* and *S4* … *Metaseiulus pomi*

19. Seta *S5* absent … subgenus *Typhlodromus* … *Typhlodromus* (*Typhlodromus*) *pyri*

Seta *S5* present … subgenus *Anthoseius* … 20

20. Dorsal shield with three pairs of solenostomes … *Typhlodromus* (*Anthoseius*) *recki*

Dorsal shield with five pairs of solenostomes … 21

21. Calyx of spermathecae short, cup, or bell-shaped … 22

Calyx of spermathecae elongated not cup, or bell-shaped … 24

22. Movable digit of chelicerae with one tooth … 23

Movable digit of chelicerae with two teeth … *Typhlodromus* (*Anthoseius*) *dossei*

23. Peritreme extending to level of seta *j1* … *Typhlodromus* (*Anthoseius*) *rhenanus*

Peritreme extending to level of seta *j3* … *Typhlodromus* (*Anthoseius*) *ueckermanni*

24. Movable digit of chelicerae with one tooth …*Typhlodromus* (*Anthoseius*) *caudiglans*

Movable digit of chelicerae with two teeth … *Typhlodromus* (*Anthoseius*) *vulgaris*

## Family Phytoseiidae Berlese

## Subfamily Amblyseiinae Muma

## Genus *Amblyseius* Berlese

### 1. Amblyseius andersoni (Chant, 1957)

#### Material examined

Netherlands, one female (HAL095F301) from *Chrysanthemum morifolium* (Asteraceae), 27 Aug 2006; Netherlands, one female (HAL095F062) from *Symphoricarpos albus* (Caprifoliaceae), 29 Oct 2006; Netherlands, two females (HAL098B508) from *Symphoricarpos* sp. (Caprifoliaceae), 20 Sept 2009; USA, two females (HAL099C254) from *Malus pumila* (Rosaceae), 25 Nov 2010; Netherlands, one female (HAL100B070) from *Skimmia* sp. (Rutaceae), 20 July 2011; Netherlands, two females (HAL101B145) from *Rosa hybrida* (Rosaceae), 20 Feb 2012; Canada, one female (QAR101H027) from *Capsicum annuum* (Solanaceae), 28 Sept 2011; Japan, one female (QAR101H019) from *Capsicum annuum* (Solanaceae), 12 Sept 2011; Netherlands, one female (QAR102H025) from *Bouvardia* sp. (Rubiaceae), 22 June 2013; Netherlands, one female (QAR102H043) from *Symphoricarpos* sp. (Caprifoliaceae), 8 Sept 2013; Netherlands, three females (QAR102H045) from *Bouvardia* sp. (Rubiaceae), 8 Sept 2013.

#### Distribution

Africa: Algeria (Athias-Henriot 1958), Morocco (Tixier *et al*. 2003). Asia: Azerbaijan (Abbasova 1972), Cyprus (Papadoulis *et al*. 2009), Japan (Chant 1959), Syria (Barbar 2013), Turkey (Çobanoğlu 1991). Europe: Austria (El-Borolossy 1989), Bosnia and Herzegovina (Döker *et al*. 2019), Czech Republic (Kabíček 2004), Denmark (Hansen & Johnsen 1986), England (Chant & Hansell 1971), France (Athias-Henriot 1962), Georgia (Wainstein & Vartapetov 1973), Germany (Westerboer & Bernhard 1963), Greece (Papadoulis & Emmanouel 1991), Hungary (Bozai 1980), Italy (Castagnoli *et al*. 1984), Lativia (Salmane 2003), Moldova (Wainstein 1973), the Netherlands (Yoshida-Shaul & Chant 1995), Poland (Boczek 1964), Portugal (Carmona 1966), Serbia (Radivojević & Petanović 1984), Slovakia (Jedlickova 1993), Slovenia (Bohinc *et al*. 2018), Spain (Villaronga & Garcia Mari 1988), Sweden (Steeghs *et al*. 1993), Switzerland (Baillod & Venturi 1980), Ukraine (Kolodochka 1973). North America: Canada (Chant 1957), the USA (Chant 1959).

#### Remarks

The species is distributed worldwide, except in Oceania and South America, and has no previous record of existing in Taiwan (Demite *et al*. 2020). EPPO (2020) listed this species as a commercially used biological control agent. Most of specimens came from the Netherlands. However, one specimen originated in Japan Japan. Toyoshima *et al*. (2016) recorded this species in Japan for the first time. Further confirmation is required to determine whether this species was introduced as a commercial product or is a native population.

### 2. Amblyseius largoensis Muma, 1955

#### Material examined

USA, one female (HAL100B062) from *Persea Americana* (Lauraceae), 19 May 2011.

#### Distribution

Africa: Angola (Carmona 1968), Benin (Zannou *et al*. 2007), Ivory Coast (Moraes *et al*. 1989a), Kenya (Swirski & Ragusa 1978), Mozambique (Rodrigues 1968), Sierra Leone (Zannou *et al*. 2007), Tanzania (El-Banhawy & Abou-Awad 1990). Asia: China (Chen *et al*. 1980), India (Gupta 1978), Iran (Daneshvar 1980), Malaysia (Ehara 2002a), Oman (Hountondji *et al*. 2010), the Philippines (Corpuz & Rimando 1966), Saudi Arabia (Alatawi *et al*. 2017), Singapore (Corpuz-Raros 1995), Sri Lanka (Moraes *et al*. 2004b), Taiwan (Ehara 1970), Thailand (Ehara & Bhandhufalck 1977), Turkey (Çobanoğlu 1989b). Central America: Cuba (Rodriguez *et al*. 1981), Dominican Republic (Ferragut *et al*. 2011), Guatemala (Chant 1959), Mexico (Chant 1959), Jamaica (Denmark & Muma 1978), Puerto Rico (De Leon 1965). Europe: Georgia (Wainstein & Vartapetov 1973). North America: USA (Muma 1955). Oceania: Australia (Collyer 1980), Fiji (Collyer 1980), Hawaii (Prasad 1968), New Caledonia (Schicha 1981b), New Zealand (Collyer 1964b), Papua New Guinea (Schicha & Gutierrez 1985), Vanuatu (Schicha 1981b). South America: Brazil (Ehara 1966), Colombia (Moraes & Mesa 1988), Guyana (De Leon 1966), Trinidad (De Leon 1967), Venezuela (Aponte & McMurtry 1993).

#### Remarks

The species is distributed worldwide; however, it is doubtful if all recorded specimens are real *A. largoensis* (e.g., Döker *et al*. 2020). Liao *et al*. (2020) indicated that the species can be identified by the parallel tubular calyx of the spermatheca. Additionally, this species is dominant species in southern Taiwan, especially in the tropical region. This species has also been studied in terms of its potential in biological control of red palm mite, *Raoiella indica* Hirst (e.g., Mendes *et al*. 2018).

### 3. Amblyseius sinuatus De Leon, 1961

#### Material examined

USA, one male (HAL095F293) from *Prunus persica* (Rosaceae), 6 July 2006; USA, one female (HAL095F299) from *Prunus persica* (Rosaceae), 24 Aug 2006; USA, two females one male (HAL099C146) from *Prunus persica* (Rosaceae), 10 Sept 2010; USA, one female (QAR101H005) from *Prunus persica* (Rosaceae), 22 June 2011; USA, one female (QAR101H003) from *Prunus persica* (Rosaceae), 4 June 2011; USA, one female two males (QAR101H004) from *Fragaria ananassa* (Rosaceae), 7 June 2011; USA, one female (QAR101H009) from *Prunus persica* (Rosaceae), 19 July 2011; USA, one female (QAR101H010) from *Prunus persica* (Rosaceae), 21 July 2011; USA, one male (QAR101H007) from *Prunus persica* (Rosaceae), 7 July 2011; USA, one female (HAL101B060) from *Prunus persica* (Rosaceae), 29 Sept 2011; USA, one female (HAL101B192) from *Prunus persica* (Rosaceae), 29 June 2012; USA, four females (QAR102H020) from *Asparagus officinalis* (Asparagaceae), 11 June 2013; USA, one female (QAR102H024) from *Prunus persica* (Rosaceae), 21 June 2013; USA, three males (QAR102H026) from *Prunus persica* (Rosaceae), 1 July 2013; USA, one male (QAR102H027) from *Prunus persica* (Rosaceae), 17 July 2013; USA, one female (QAR102H028) from *Prunus persica* (Rosaceae), 17 July 2013.

#### Distribution

North America: Mexico (De Leon 1961).

#### Remarks

This species is distributed only in Mexico (De Leon 1961; Demite *et al*. 2020); however, 17 samples were collected from the United States; the distribution of the species requires further confirmation. Additionally, most of the specimens were were obtained from *Prunus persica*.

### 4. Amblyseius tamatavensis Blommers, 1974

#### Material examined

Thailand, one female one male (HAL098B608) from *Piper nigrum* (Piperaceae), 30 Nov 2009; Thailand, one female (HAL099B097) from *Aranthera beatrice* (Orchidaceae), 6 Jan 2010; Thailand, one male (HAL100B201) from *Piper betle* (Piperaceae), 27 Oct 2011; Thailand, one female (HAL101B018) from *Piper betle* (Piperaceae), 6 Dec 2011; Malaysia, one female (HAL101B189) from *Ocimum tenuiflorum* (Lamiaceae), 16 June 2012.

#### Distribution

Africa Benin (Zannou *et al*. 2007), Burundi (Zannou *et al*. 2007), Cameroon (Zannou *et al*. 2007), Dr Congo (Zannou *et al*. 2007), Ghana (Zannou *et al*. 2007), Kenya (Moraes *et al*. 1989b), Madagascar Island (Blommers 1974), Malawi (Zannou *et al*. 2005), Mozambique (Zannou *et al*. 2005), Nigeria (Moraes *et al*. 1989a), Rwanda (Zannou *et al*. 2007), South Africa (Ueckermann & Loots 1988), Uganda (Zannou *et al*. 2007). Asia: Indonesia (Oomen 1982), Japan (Ehara & Amano 2002), Malaysia (Ehara 2002a), the Philippines (Schicha & Corpuz-Raros 1992), Singapore (Corpuz-Raros 1995), Sri Lanka (Moraes *et al*. 2004b), Thailand (Oliveira *et al*. 2012), Taiwan (Liao *et al*. 2013). Central America: Cuba (Moraes *et al*. 1991), Dominican Republic (Abo-Shnaf *et al*. 2016). North America: USA (Döker *et al*. 2018). Oceania: Australia (Schicha 1981a), Fiji (Gutierrez & Schicha 1984), Papua New Guinea (Schicha 1981a), Vanuatu (Schicha 1981a), Western Samoa (Schicha 1981a). South America: Brazill (Gondim Jr. & Moraes 2001), Venezuela (Quirós *et al*. 2005).

#### Remarks

Liao *et al*. (2020) provided a detailed redescription of the species and considered the tooth numbers of movable and fixed digits as important characters. Döker *et al*. (2018) reported the biological control potential of this species on whiteflies, and Ho & Chen (2001) also reported that the species has the potential to become an effective predator for *Thrips palmi* Karny.

### 5. Amblyseius tsugawai Ehara, 1959

#### Material examined

Japan, one female (QAR101H028) from *Perilla frutescens* (Lamiaceae), 28 Sept 2011.

#### Distribution

Asia: China (Zhu & Chen 1983), Japan (Ehara 1959), South Korea (Ryu & Kim 1998).

#### Remarks

This species is distributed in East Asia, especially Japan, and is a major predator of spider mites in orchards in Japan. It is considered as Type III generalist predators (Funayama & Sonoda 2014). The food sources of the species ranges from spider mites, thrips, whiteflies, pyralid moths, and pollens (Yang *et al*. 2019).

## Genus *Neoseiulus* Hughes

### 6. Neoseiulus barkeri Hughes, 1948

#### Material examined

Thailand, one female (HAL095F297) from *Asparagus officinalis* (Asparagaceae), 15 Aug 2006; Thailand, one female (HAL095F298) from *Ocimum sanctum* (Lamiaceae), 21 Aug 2006; Netherlands, one female (HAL095F309) from *Lactuca sativa var. capitata* (Asteraceae), 14 Sept 2006; Israel, one female (HAL095G063) from *Kochia* sp. (Amaranthaceae), 31 Oct 2006; Israel, three females (HAL095G064) from *Limonium* sp. (Plumbaginaceae), 31 Oct 2006; Japan, one female (HAL100B075) from *Vitis vinifera* (Vitaceae), 2 May 2011; Thailand, one female (HAL101B144) from *Asparagus officinalis* (Asparagaceae), 13 Feb 2012; New Zealand, one female (QAR101H001) from *Malus pumila Yoko* (Rosaceae), 5 Apr 2011; Japan, one female (QAR101H013) from *Perilla frutescens* (Lamiaceae), 23 Aug 2011; USA, one female (QAR101H014) from *Anthriscus cerefolium* (Apiaceae), 27 Aug 2011; Thailand, one protonymph (QAR101H015) from *Asparagus officinalis* (Asparagaceae), 30 Aug 2011; Japan, one female (QAR101H023) from *Chrysanthemum* sp. (Asteraceae), 21 Sept 2011; Thailand, one female (QAR101H022) from *Asparagus officinalis* (Asparagaceae), 17 June 2013; Peru, six females (QAR101H029) from *Asparagus officinalis* (Asparagaceae), 21 July 2013; Japan, one female (QAR101H036) from *Perilla frutescens* (Lamiaceae), 3 Sept 2013; Netherlands, one female (QAR101H038) from *Thymes* sp. (Lamiaceae), 5 Sept 2013; Japan, one female (QAR101H050) from *Perilla frutescens* (Lamiaceae), 13 Sept 2013.

#### Distribution

Africa: Algeria (Athias-Henriot 1961), Benin (Zannou *et al*. 2006), Burundi (Zannou *et al*. 2006), Canary Islands (Ferragut & Peña-Estévez 2003), Cape Verde (Ueckermann 1992), Egypt (Abo-Shnaf & Moraes 2014), Ghana (Moraes *et al*. 1989a), Guinea (Ragusa & Athias-Henriot 1983), Kenya (Zannou *et al*. 2006), Malawi (Zannou *et al*. 2005), Morocco (Kreiter *et al*. 2004), Mozambique (Zannou *et al*. 2006), Nigeria (Moraes *et al*. 1989a), Senegal (Kade *et al*. 2011), South Africa (Ueckermann & Meyer 1988), Tunisia (Kreiter *et al*. 2002), Yemen (Ueckermann 1996). Asia: China (Wu 1981), Cyprus (Amitai 1992), Iran (Rahmani *et al*. 2010), Israel (Swirski & Amitai 1965), Japan (Ehara 1972), Jordan (Allawi 1991), Oman (Hountondji *et al*. 2010), Saudi Arabia (Al-Atawi 2011b), South Korea (Ryu 1997), Syria (Barbar 2013), Thailand (Oliveira *et al*. 2012), Turkey (Swirski & Amitai 1982). Europe: England (Hughes 1948), Finland (Tuovinen 1993), France (Kreiter *et al*. 2000), Georgia (Wainstein & Vartapetov 1973), Germany (Karg 1965), Greece (Papaioannou-Souliotis 1981), Italy (Athias-Henriot 1961), Latvia (Petrova *et al*. 2000), the Netherlands (van de Vrie 1963), Norway (Denmark & Edland 2002), Portugal (Ferreira & Carmona 1994), Russia (Meshkov 1999), Spain (Ragusa & Athias-Henriot 1983), Sweden (Steeghs *et al*. 1993), Ukraine (Wainstein & Shcherbak 1972). North America: USA (Denmark & Evans 2011). Oceania: Australia (Waite & Gerson 1994), Hawaii (Denmark & Evans 2011). South America: Brazil (Moraes *et al*. 1993), Chile (Ragusa & Vargas 2002).

#### Remarks

Beaulieu & Beard (2018) provided a detailed redescription and illustration of this species. They also designated a neotype of the species. The species is distributed worldwide, including in Taiwan (Demite *et al*. 2020; Liao *et al*. 2020). The species is considered a subtype III-e generalist predators that lives on soil/litter, and plants (McMurtry *et al*. 2013). EPPO (2020) listed this species as a commercial biological control agent for spider mites and thrips.

### 7. *Neoseiulus bicaudus* (Wainstein, 1962)

#### Material examined

Japan, one female (HAL095F295) from *Vitis vinifera* (Vitaceae), 3 Aug 2006; Japan, one female (HAL095F308) from *Vitis vinifera* (Vitaceae), 8 Sept 2006.

#### Distribution

Africa: Egypt (Abo-Shnaf & Moraes 2014), Tunisia (Sahraoui *et al*. 2012). Asia: Armenia (Arutunjan 1969), Azerbaijan (Abbasova 1972), Iran (Asali Fayaz *et al*. 2011), Israel (Swirski & Amitai 1985), Kazakhstan (Wainstein 1962), Saudi Arabia (Negm *et al*. 2012), Syria (Barbar 2014), Tajikistan (Wainstein 1962), Turkey (Çobanoğlu 1991). Europe: France (Athias-Henriot 1966), Georgia (Wainstein & Vartapetov 1973), Greece (Papadoulis & Emmanouel 1990), Hungary (Bozai 1980), Italy (Ragusa & Paoletti 1985), Latvia (Salmane 1996), Moldova (Kolodochka 1980), Norway (Denmark & Edland 2002), Portugal (Espinha *et al*. 1998), Russia (Wainstein 1962), Serbia (Stojnić *et al*. 2002), Slovakia (Fend’a 2010), Spain (Iraola *et al*. 1997), Switzerland (Airoldi *et al*. 1989), Ukraine (Livshitz & Kuznetsov 1972). North America: Mexico (Denmark & Evans 2011), USA (Congdon 2002). South America: Chile (Trincado *et al*. 2018).

#### Remarks

The species is distributed worldwide, except in Oceania. It is a native predator of spider mites and thrips in Xinjiang Uygur Autonomous Region of China and is more adapted than nonnative species to hot and dry climates (Zhang *et al*. 2017).

### 8. Neoseiulus californicus (McGregor, 1954)

#### Material examined

Japan, three females one male (HAL095F372) from *Vitis vinifera* (Vitaceae), 26 Aug 2006; Japan, one female (HAL095F307) from *Vitis vinifera* (Vitaceae), 31 Aug 2006; USA, four females (HAL099C147) from *Hydrangea macrophylla* (Hydrangeaceae), 2 Oct 2010; Chile, three females (HAL100B069) from *Malus pumila* (Rosaceae), 1 May 2011; Chile, three females (HAL100B073) from *Malus pumila* (Rosaceae), 1 May 2011; USA, two females (QAR101H033) from *Fragaria ananassa* (Rosaceae), 12 Sept 2011; USA, two females (QAR101H035) from *Fragaria ananassa* (Rosaceae), 19 Sept 2011; USA, one female (QAR101H030) from *Thymes* sp. (Lamiaceae), 18 Oct 2011; Japan, one female (HAL101B180) from *Perilla frutescens* (Lamiaceae), 30 Mar 2012; Chile, one female (HAL101B186) from *Malus pumila* (Rosaceae), 30 May 2012; Chile, one female (HAL101B188) from *Malus pumila* (Rosaceae), 8 June 2012; USA, one female (QAR102H031) from *Prunus persica* (Rosaceae), 7 July 2013.

#### Distribution

Africa: Morocco (Tixier *et al*. 2016), Senegal (Kade *et al*. 2011), South Africa (Villiers & Pringle 2011), Tunisia (Kreiter *et al*. 2002). Asia: Cyprus (Vassiliou *et al*. 2012), Japan (Amano 1994), South Korea (Jung *et al*. 2006), Syria (Barbar 2014), Turkey (Çakmak & Çobanoğlu 2006), Vietnam (Nguyen *et al*. 2019). Central America: Guatemala (McMurtry 1977). Europe: France (Athias-Henriot 1977), Greece (Papaioannou-Souliotis *et al*. 1999), Italy (Vacante & Nucifora 1987), Portugal (Ferreira & Carmona 1994), Serbia (Stojnić *et al*. 2002), Slovenia (Bohinc *et al*. 2018), Spain (McMurtry 1977). North America: Canada (Denmark & Evans 2011), Cuba (Ramirez *et al*. 1988), Mexico (Estebanes-Gonzale & Rodriguez-Navarro 1998), USA (McGregor 1954). South America: Argentina (McMurtry 1977), Brazil (Ferla & Moraes 1998), Chile (Athias-Henriot 1977), Colombia (Moraes & Mesa 1988), Peru (McMurtry 1977), Venezuela (Aponte & McMurtry 1993).

#### Remarks

McGregor (1954) described the species based on male specimen. Numerous acarologists later provided opinions on the species. Beaulieu & Beard (2018) provided a detailed redescription and illustration and designated a neotype of the species. They maintained the species name “*N. californicus*” from the species concept of Athias-Henriot (1977), which is its prevailing usage as commercial products for biological control.

Although the species has characteristics of a Type III generalist predator due to its wider range of food sources (e.g., spider mites, tarsonemid mites, thrips, and pollens), it is almost always associated with tetranychids that produce heavy webbing. Therefore, it is still classified as Type II selective predators of tetranychids (McMurtry *et al*. 2013). Moreover, Döker *et al*. (2016) reported that *N. californicus* can survive in extremely humid conditions. EPPO (2020) listed this species as a commercial biological control agent for spider mites and thrips.

### 9. Neoseiulus cucumeris (Oudemans, 1930)

#### Material examined

Australia, one female (HAL098B609) from *Hydrangea macrophylla* (Hydrangeaceae), 28 Oct 2009; Netherlands, one female (HAL101B179) from *Helleborus* sp. (Ranunculaceae), 30 Mar 2012; Netherlands, one female (QAR101H008) from *Trachelium caeruleum* (Campanulaceae), 18 July 2011; Netherlands, one female (QAR101H011) from *Hydrangea macrophylla* (Hydrangeaceae), 18 Aug 2011; Netherlands, one female (QAR102H021) from *Hydrangea macrophylla* (Hydrangeaceae), 16 June 2013; Netherlands, one female (QAR102H051) from *Hydrangea macrophylla* (Hydrangeaceae), 15 Sept 2013.

#### Distribution

Africa: Algeria (Athias-Henriot 1960), Egypt (Chant 1959), Morocco (McMurtry & Bounfour 1989), Tunisia (Kreiter *et al*. 2004). Asia: Armenia (Arutunjan 1970), Azerbaijan (Gadzhiev & Abbasova 1965), Cyprus (Vassiliou *et al*. 2012), India (Sadana & Kanta 1971), Iran (Sepasgozarian 1977), Israel (Amitai & Swirski 1978), Saudi Arabia (Al-Atawi 2011a), Turkey (Özman & Çobanoğlu 2001). Europe: Austria (Bohm 1960), Belarus (Sidlyarevich 1966), England (Collyer 1956), Finland (Tuovinen 1993), France (Oudemans 1930b), Georgia (Wainstein 1961), Germany (Dosse 1956), Greece (Papadoulis & Emmanouel 1991), Hungary (Bozai 1980), Italy (Ragusa 1977), Latvia (Petrova *et al*. 2000), Moldova (Wainstein 1973), Netherlands (Chant 1959), Norway (Denmark & Edland 2002), Poland (Wiackowski & Suski 1963), Portugal (Espinha *et al*. 1998), Russia (Meshkov 1999), Slovakia (Fend’a & Schniererová 2005), Slovenia (Bohinc & Trdan 2013), Spain (Escudero & Ferragut 1998), Sweden (Sellnick 1958), Switzerland (Chant 1959), Ukraine (Livshitz & Kuznetsov 1972). North America: Canada (Nesbitt 1951), Mexico (Chant 1959), USA (Nesbitt 1951). Oceania: Australia (Beard 2001), New Zealand (Chant 1959). South America: Chile (Ragusa & Vargas 2002).

#### Remarks

The species is distributed worldwide, but had no previous record in Taiwan (Demite *et al*. 2020). EPPO (2020) listed this species as a commercially used biological control agent. This species has the lifestyle of Type III-e, generalist predator, which has soil/litter habitats similar to *N. barkeri*. Both species considered effective natural enemies of thrips and spider mites.

### 10. *Neoseiulus imbricatus* (Corpuz & Rimando, 1966)

#### Material examined

Thailand, one female (HAL100B196) from *Asparagus officinalis* (Asparagaceae), 21 Nov 2011.

#### Distribution

Asia: Azerbaijan (Abbasova 1972), China (Zhu & Chen 1983), India (Gupta 1986), Philippines (Corpuz & Rimando 1966), Saudi Arabia (Alatawi *et al*. 2017), Thailand (Ehara & Bhandhufalck 1977).

#### Remarks

This species is distributed in Asia (Demite *et al*. 2020). No difference exists between our specimens and the paratype specimen (Aca014 from UPLB-MNH). Wu *et al*. (2009) reported that the species is an effective natural enemy of spider mites and tarsonemids in rice fields.

### 11. *Neoseiulus loxus* (Schuster & Pritchard, 1963)

#### Material examined

USA, one female (HAL095F305) from *Fragaria chiloensis* (Rosaceae), 29 Aug 2006; USA, one female (HAL095F306) from *Fragaria chiloensis* (Rosaceae), 29 Aug 2006; USA, one female (HAL095F315) from *Fragaria chiloensis* (Rosaceae), 3 Oct 2006.

#### Distribution

North America: USA (Schuster & Pritchard 1963).

#### Remarks

The species is only distributed in the United States (Demite *et al*. 2020). Schuster & Pritchard (1963) reported the species from *Zostera marina* on sea shore, a species habitat for phytoseiid mites. Denmark & Evans (2011) recorded additional habitat plants, including strawberry, and *Malus* sp. The collecting records of the species are all from *Fragaria chiloensis.* The biological control potential of the species requires further confirmation.

### 12. Neoseiulus makuwa (Ehara, 1972)

#### Material examined

Japan, one female (HAL095F296) from *Vitis vinifera* (Vitaceae), 3 Aug 2006; Japan, one female (HAL095F310) from *Brassica rapa* (Brassicaceae), 15 Sept 2006; Thailand, one female (QAR102H022) from *Lactuca sativa* (Asteraceae), 20 June 2013.

#### Distribution

Africa: Cameroon (Zannou *et al*. 2006). Asia: China (Zhu & Chen 1983), Indonesia (Ehara 2002b), Japan (Ehara 1972), Saudi Arabia (Negm *et al*. 2012), South Korea (Ryu 1993), Taiwan (Tseng 1983), United Arab Emirates (Negm 2014).

#### Remarks

The species is distributed in Africa and Asia (Demite *et al*. 2020). Ehara (1972) described this species from melons in Japan. No difference was observed among these specimens and holotype (NSMT-AC13110 from NSMT). Liao *et al*. (2020) reported a correlation between the species and tetranychid mites, but further experiments are required to determine its biological control potential.

## Genus Phytoseiulus Evans

### 13. *Phytoseiulus persimilis* Athias-Henriot, 1957

#### Material examined

Netherlands, three females one male (HAL095F302) from *Symphoricarpos albus* (Caprifoliaceae), 27 Aug 2006; Netherlands, three females one male (HAL095F303) from *Symphoricarpos albus* (Caprifoliaceae), 27 Aug 2006; Netherlands, one female (HAL095F317) from *Lactuca sativa var. capitata* (Asteraceae), 12 Oct 2006; New Zealand, three females (HAL099C250) from *Fragaria ananassa* (Rosaceae), 29 Nov 2010; USA, one male (QAR101H016) from *Fragaria ananassa* (Rosaceae), 30 Aug 2011.

#### Distribution

Africa: Algeria (Athias-Henriot 1957a), Egypt (Afsah 2015), Kenya (Migeon *et al*. 2019), Lybia (Damiano 1961), Mauritius (Kreiter *et al*. 2018), Morocco (McMurtry & Bounfour 1989), South Africa (Meyer 1981), Tunisia (Rambier 1972). Asia: China (Wu *et al*. 1997), Cyprus (Vassiliou *et al*. 2012), Iran (Hajizadeh & Mortazavi 2015), Israel (Swirski & Amitai 1968), Japan (Ohno *et al*. 2012), Jordan (Allawi 1991), Philippine (Corpuz-Raros 2005), Syria (Barbar 2013), Turkey (Sekeroglu & Kazak 1993). Central America: Costa Rica (Denmark *et al*. 1999), Guatemala (Denmark *et al*. 1999). Europe: Finland (Tuovinen 1993), France (Rambier 1972), Greece (Swirski & Ragusa 1976), Hungary (Bozai 1996), Italy (Kennett & Caltagirone 1968), Latvia (Salmane 2001), Portugal (Ferreira & Carmona 1994), Serbia (Kropczynska & Petanović 1987), Slovenia (Kreiter *et al*. 2020), Spain (Ferragut *et al*. 1983). North America: USA (Denmark & Evans 2011). Oceania: Australia (Goodwin & Schicha 1979). South America: Chile (Gonzalez 1961), Peru (El-Banhawy 1979), Venezuela (Aponte & McMurtry 1993).

#### Remarks

The species is distributed worldwide but had no previous record in Taiwan (Demite *et al*. 2020). Lo *et al*. (1986) reported that the species was introduced for biological purposes from the United States. EPPO (2020) listed this species as a commercially used biological control agent of Type I-a, specialized predators of *Tetranychus* spider mites (McMurtry *et al*. 2013).

## Genus Proprioseiopsis Muma

### 14. Proprioseiopsis asetus (Chant, 1959)

#### Material examined

Japan, one female (HAL095F311) from *Brassica rapa* (Brassicaceae), 15 Sept 2006.

#### Distribution

Asia: China (Wu *et al*. 2009), Saudi Arabia (Negm *et al*. 2012), Taiwan (Tseng 1983), Unite Arab Emirates (Negm 2014). Central America: Cuba (Ramos & Rodriguez 1999), Jamaica (Denmark & Muma 1978), Nicaragua (Rodríguez-Morell *et al*. 2013). North America: Mexico (Estebanes-Gonzale & Rodriguez-Navarro 1998), USA (Chant 1959). Oceania: Hawaii (Wainstein 1983). South America: Brazil (Denmark & Muma 1973).

#### Remarks

Liao *et al*. (2020) provided a detailed comparison between the species and *Prop. mexicanus*. Although no significant morphological differences were found, the study disagrees with the synonymy proposed by Denmark & Evans (2011) due the different biology. Two hypotheses require further study: (1) *Prop. mexicanus* (in the United States) and *Prop. asetus* (in China and Taiwan) are two different species; (2) the mites have different life histories in different locations (parthenogenesis in China and Taiwan).

The species is considered as lifestyle Type III-e, generalist predators from soil/litter habitats. Ho & Chen (2001) proposed that the mites have the potential to become natural enemies of thrips.

### 15. Proprioseiopsis neotropicus (Ehara, 1966)

#### Material examined

Peru, one female (QAR102H029) from *Asparagus officinalis* (Asparagaceae), 21 July 2013.

#### Distribution

South America: Argentina (Guanilo *et al*. 2008a), Brazil (Ferla & Moraes 1998), Colombia (Moraes & Mesa 1988), Ecuador (Moraes *et al*. 1991), Peru (Guanilo *et al*. 2008b).

#### Remarks

The species is distributed in South America (Demite *et al*. 2020). It is a generalist predator inhabiting natural vegetation and crop fields (Moraes *et al*. 2004a). The biological control potential of the species requires further exploration.

## Subfamily Typhlodrominae Scheuten

## Genus Galendromus Muma

## Subgenus *Galendromus* Muma

### 16. Galendromus (Galendromus) occidentalis (Nesbitt, 1951)

#### Material examined

USA, one female (HAL095F371) from *Prunus persica* (Rosaceae), 25 Sept 2006; USA, six females (HAL095F350) from *Malus pumila* (Rosaceae), 9 Nov 2006; USA, one female (HAL095F455) from *Malus pumila* (Rosaceae), 15 Nov 2006; USA, two females (HAL095F456) from *Malus pumila* (Rosaceae), 15 Nov 2006; USA, four females (HAL095F458) from *Malus pumila* (Rosaceae), 15 Nov 2006; USA, two females (HAL095F459) from *Malus pumila* (Rosaceae), 15 Nov 2006; USA, two females (TAL095F457) from *Prunus persica* (Rosaceae), 15 Nov 2006; USA, four females (HAL095G068) from *Malus pumila* (Rosaceae), 30 Nov 2006; USA, four females (HAL095G069) from *Malus pumila* (Rosaceae), 30 Nov 2006; USA, three females (HAL095G070) from *Malus pumila* (Rosaceae), 1 Dec 2006; USA, seven females (HAL095G079) from *Malus pumila* (Rosaceae), 7 Dec 2006; USA, four females (HAL095G081) from *Malus pumila* (Rosaceae), 7 Dec 2006; USA, seven females (HAL095G799) from *Malus pumila* (Rosaceae), 19 Dec 2006; USA, nine females (HAL095G800) from *Malus pumila* (Rosaceae), 25 Dec 2006; USA, three females (QAR101H022) from *Prunus persica* (Rosaceae), 19 Sept 2011; USA, one female (HAL101B056) from *Prunus persica* (Rosaceae), 21 Sept 2011; USA, one female (QAR101H024) from *Prunus persica* (Rosaceae), 22 Sept 2011; USA, one female (QAR101H025) from *Prunus persica* (Rosaceae), 22 Sept 2011; USA, one female (QAR101H026) from *Prunus persica* (Rosaceae), 22 Sept 2011; USA, three females (HAL101B059) from *Prunus persica* (Rosaceae), 28 Sept 2011; USA, one female (QAR102H034) from *Prunus persica* (Rosaceae), 14 Aug 2013; USA, one female (QAR102H046) from *Prunus persica* (Rosaceae), 12 Sept 2013; USA, one female (QAR102H047) from *Prunus persica* (Rosaceae), 12 Sept 2013; USA, one female (QAR102H053) from *Prunus persica* (Rosaceae), 26 Sept 2013.

#### Distribution

Africa: South Africa (Denmark 1982). Asia: China (Wu *et al*. 1997), Israel (Denmark 1982), Jordan (Allawi 1991), South Korea (Ryu & Lee 1992), Taiwan (Chant & Yoshida-Shaul 1984). Europe: Austria (El-Borolossy 1989), Greece (Ragusa Di Chiara *et al*. 1995), Netherlands (Hoying & Croft 1977), Russia (Denmark 1982). North America: Canada (Nesbitt 1951), Mexico (Denmark & Evans 2011), USA (Cunliffe & Baker 1953). Oceania: Australia (Whitney & James 1996), New Zealand (Collyer 1964b). South America: Chile (Prado 1991), Venezuela (Aponte & McMurtry 1993).

#### Remarks

The species is distributed worldwide (Demite *et al*. 2020). Denmark (1982) reported species distribution in Taiwan without recording specimen information. Chant & Yoshida-Shaul (1984) reported the species to be distributed in Taiwan based on one female specimen collected in 1950. Lo *et al*. (1986) reported that the species was introduced to Taiwan but could not be kept in laboratory. All specimens were collected on peaches and apples from the United States. The species has favorable efficacy for use in orchards. EPPO (2020) listed this species as a commercially used biological control agent.

## Genus *Meyerius* van der Merwe

### 17. *Meyerius immutatus* (van der Merwe, 1968)

#### Material examined

South Africa, one female (HAL095G059) from *Berzelia* sp. (Bruniaceae), 27 Oct 2006; South Africa, one female (HAL095G060) from *Berzelia* sp. (Bruniaceae), 27 Oct 2006; South Africa, one female (HAL095G051) from *Brunia laevis* (Bruniaceae), 28 Oct 2006; South Africa, one male (HAL095G059) from *Berzelia* sp. (Bruniaceae), 1 Dec 2006.

#### Distribution

Africa: South Africa (Van der Merwe 1968).

#### Remarks

The species has only been recorded in South Africa (Demite *et al*. 2020). All species were found from commodity of South Africa in 2006, and it should be noted that the related habitat plants were only imported during this time.

## Genus Metaseiulus Muma

## Subgenus *Metaseiulus* Chant and McMurtry

### 18. Metaseiulus (Metaseiulus) pomi (Parrott, 1906)

#### Material examined

Thailand, two females one male (HAL100B063) from *Rubus idaeus* (Rosaceae), 30 June 2011.

#### Distribution

Europe: the Netherlands (Faraji 2006). North America: Canada (Nesbitt 1951), USA (Parrott *et al*. 1906).

#### Remarks

The species has only been recorded in the Netherlands, Canada, and the United States (Demite *et al*. 2020). The specimens were collected from commodity of Thailand, which had no previous record of the species. The distribution of this species requires further confirmation.

## Genus *Typhlodromus* Scheuten

## Subgenus *Anthoseius* De Leon

### 19. Typhlodromus (Anthoseius) caudiglans Schuster, 1959

#### Material examined

USA, one female (HAL095F312) from *Prunus persica* (Rosaceae), 16 Sept 2006; USA, one female (QAR101H020) from *Prunus persica* (Rosaceae), 16 Sept 2011; USA, one female (QAR101H021) from *Prunus persica* (Rosaceae), 19 Sept 2011; USA, one female (HAL101B058) from *Prunus persica* (Rosaceae), 23 Sept 2011; USA, one female (QAR102H019) from *Prunus persica* (Rosaceae), 8 June 2013; USA, one female (QAR102H047) from *Prunus persica* (Rosaceae), 12 Sept 2013; Japan, three females (QAR102H049) from *Vitis vinifera* (Vitaceae), 13 Sept 2013; USA, onefemale (QAR102H052) from *Prunus persica* (Rosaceae), 17 Sept 2013.

#### Distribution

Asia: Azerbaijan (Abbasova 1972), China (Wu 1988), Iran (Hajizadeh *et al*. 2002), Europe: Austria (El-Borolossy 1989), England (Collyer 1964), Latvia (Salmane & Petrova 2002), Lithuania (Pauriene 1970), Moldova (Beglyarov & Malov 1977), Norway (Evans & Edland 1998), Russia (Wainstein 1975), Slovakia (Jedlickova & Kolodochka 1994), Ukraine (Kolodochka 1978). North America: Canada (Downing & Moilliet 1971), USA (Schuster 1959). Oceania: Australia (Chant *et al*. 1978), New Zealand (Collyer 1964a).

#### Remarks

The species is distributed worldwide, except in Africa and South America (Demite *et al*. 2020). Most of the specimens were found on peaches from the United States, but one was found on grapes from Japan. The species may become a natural enemy in orchards.

### 20. *Typhlodromus* (*Anthoseius*) *ueckermanni* Liao & Ho sp. nov. (Figures 1–8)

#### Diagnosis

Female dorsal surface strongly reticulated, bearing 20 pairs of dorsal setae (including *r3*, *R1*). All setae smooth except *Z5* which is slightly serrated. Five pairs of solenostomes, (*gd2*, *gd4*, *gd6*, *gd8*, *gd9*) visible on the dorsal shield. Peritreme extending to level of seta *j3*. Sternal shield with two pairs of setae; ventrianal shield bearing four pairs of pre-anal setae, with a small rounded solenostomes. Fixed digit of chelicera with four teeth; movable digit with one tooth. Calyx of spermatheca cup-shaped with distal half lightly sclerotized. Leg IV with one pair of macrosetae; genu II with eight setae.

**Female (n=4)**

A lightly sclerotized mite. Idiosomal setal pattern: 12A:8A/JV:ZV.

##### *Dorsum* (Fig. 1)

Dorsal shield nearly oval, constricted at level of *R1*, strongly reticulated; **333** 329 (314–339) long (*j1*–*J5* level) and **176** 169 (159–176) wide at level of *j6*, **182** 174 (163–182) wide at level of *S4*; five pairs of solenostomes on dorsal shield, (*gd2*, *gd4*, *gd6*, *gd8*, *gd9*), thirteen pairs of lyrifissures, (*id1*, *id2*, *id4*, *id6*, *idm2*, *idm3*, *idm4*, *idm5*, *idm6*, *is1*, *idl2*, *idl3*, *idl4*); length of setae: *j1* **21** 21 (20–22), *j3* **24** 27 (24–29), *j4* **15** 17 (15–20), *j5* **15** 16 (15–17), *j6* **17** 20 (17–23), *J2* **20** 22 (20–25), *J5* **11** 9 (6–11), *z2* **18** 20 (18–22), *z3* **28** 27 (26–28), *z4* **22** 25 (22–26), *z5* **17** 19 (17–22), *Z4* **34** 33 (31–35), *Z5* **58** 57 (54–59), *s4* **27** 30 (27–33), *s6* **27** 32 (27–35), *S2* **30** 32 (30–34), *S4* **28** 30 (28–31), *S5* **27** 26 (25–27), *r3* **33** 31 (28–33), *R1* **23** 26 (23–29). All setae smooth, sharp pointed, except for *Z5* slightly serrated.

##### *Peritreme* (Figs. 1, 2)

Peritreme extending to level of seta *j3*; peritremal shield lightly sclerotized, with one pair of solenostomes (*gd3*) and one pair of lyrifissures (*id3*).

##### *Venter* (Fig. 2)

Sternal shield smooth, posterior margin irregular, much wider than long, **54** 53 (49–58) long, **62** 66 (62–71) wide, with two pairs of setae *st1* **27** 25 (22–27), *st2* **27** 24 (20–27), two pairs of lyrifissures (*pst1*, *pst2*). Sternal seta *st3* **23** 23 (21–24) on separated platelets. Exopodal shield at coxae II–IV. Metasternal platelets tear-shaped, with one pair of metasternal setae, *st4* **25** 26 (25–28), with one pair of lyrifissures (*pst3*). Genital shield smooth, truncate posteriorly, with one pair of genital setae *st5* **22** 21 (18–22), **60** 61 (57–66) wide at level of genital setae. Distances between *st1*–*st1* **51** 51 (48–52), *st2*–*st2* **51** 55 (51–58), *st5*–*st5* **58** 54 (50–58). Ventrianal shield pentagonal with waist at *JV2* level, reticulated, **104** 104 (96–115) long, **83** 81 (79–83) wide at level of *ZV2*, **73** 74 (71–77) wide at level of anus; with four pairs of pre-anal setae, *JV1* **17** 17 (17–19), *JV2* **15** 13 (10–15), *JV3* **12** 15 (12–20), *ZV2* **13** 16 (13–19), solenostomes *gv3* small and rounded; *Pa* **13** 12 (11–13), *Pst* **11** 12 (11–15) on shield. Setae *JV4* **18** 16 (13–19), *JV5* **53** 49 (47–53), *ZV1* **17 1**8 (14–21), *ZV3* **9** 12 (9–14) on interscutal membrane. All setae smooth, sharp pointed. Two metapodal plates **25** 27 (25–29) long, **4** 4 (3–4) wide, **11** 10 (11–13) long, **1** 2 (1–2) wide.

##### Spermatheca (Fig. 4)

Calyx bell-shaped and elongated, flaring distally, with apical half thick (sclerotized), **17** 18 (17–20) long, **9** 11 (9–13) wide, atrium incorporated with the calyx, major duct broad.

##### Chelicera (Fig. 3)

Movable digit **26** 27 (26–27) long, with one tooth; fixed digit **25** 26 (25–27) long, with four teeth, and with pilus dentilis.

##### *Legs* (Figs. 5–8)

Complement of setae on coxae I-IV: 2-2-2-1. Chaetotaxy (femur to basitarsus): leg I, 2-3/1-2/2-2, 2-2/1-1/1-2, 2-2/1-2/1-2, 1-1/0-1/0-1; leg II, 2-3/1-2/1-1, 2-2/1-2/0-1, 1-1/1-2/1-1, 1-1/0-1/0-1; leg III, 1-2/1-1/0-1, 1-2/1-2/0-1, 1-1/1-2/1-1, 1-1/0-1/0-1; leg IV, 1-2/1-1/0-1, 1-2/1-2/0-1, 1-1/1-2/0-1, 1-1/0-1/0-1. Macrosetae: *St* IV (pd) **33** 30 (28–33). Marcrosetae setiform.

**FIGURES 1–4.**
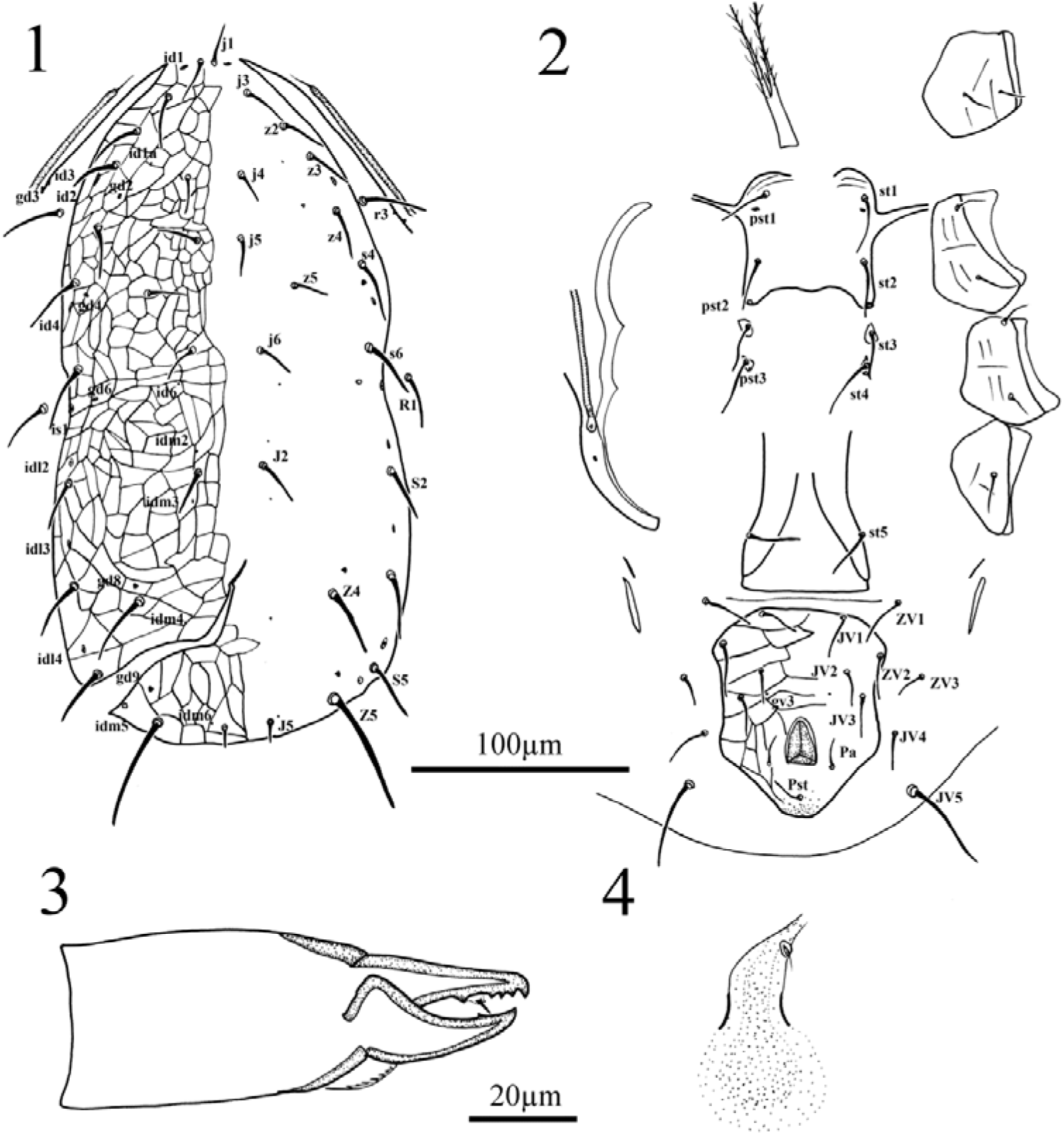
*Typhlodromus* (*Anthoseius*) *ueckermanni* **sp. nov.,** Female. 1. dorsal shield; 2. ventral idiosoma; 3. chelicera; 4. spermatheca.

**FIGURES 5–8.**
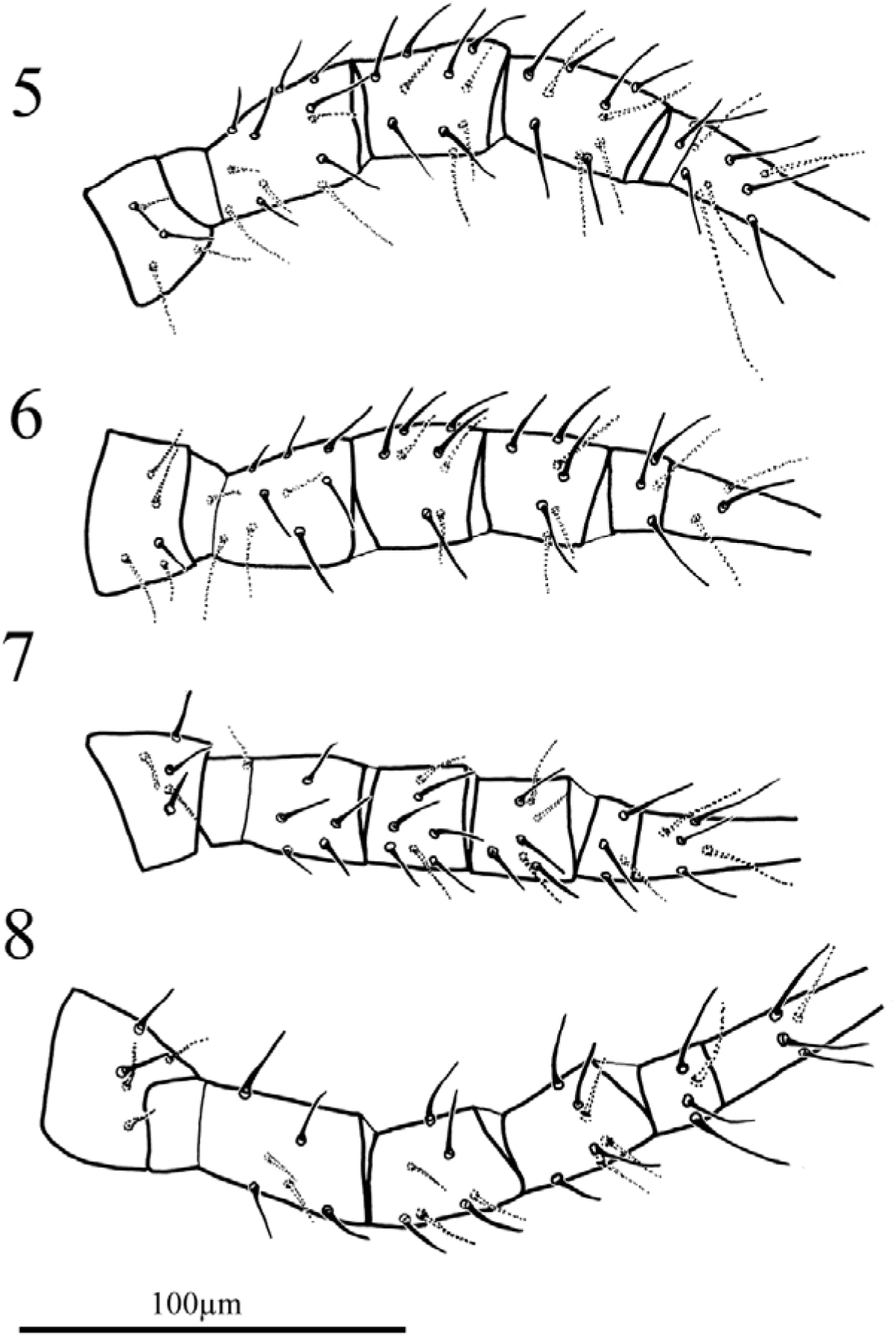
*Typhlodromus* (*Anthoseius*) *ueckermanni* **sp. nov.**, Female, legs. 5. leg I posterior view; 6. leg II posterior view; 7. leg III anterodorsal view; 8. leg IV anterior view.

#### Type specimens. Female

Holotype: South Africa, one female (HAL095G078) from *Rosmarinus officinalis* (Lamiaceae), 7 Dec 2006. Paratypes: three females (HAL095G078) data same with holotype.

#### Etymology

The epithet “*ueckermanni*“ refers to famous Acarologist from South Africa- Prof. Eddie Ueckermann, who provided detailed comparison with this species and *T.* (*A.*) *lootsi*.

#### Distribution

Asia: South Africa (present study).

#### Remarks

The new species was compared to all known species of the subgenus *Anthoseius*. Among them, it shows a close affinity to *T*. (*A*.) *aktherecus* (Kolodochka), *T*. (*A*.) *bagdasarjani* Wainstein & Arutunjan, *T*. (*A*.) *georgicus* Wainstein, *T*. (*A*.) *involutus* Livshitz & Kuznetsov, *T.* (*A.*) *karaisaliensis* Döker & Kazakkerkirae, *T.* (*A.*) *lootsi* Schultz, *T*. (*A*.) *ponticus* (Kolodochka), *T*. (*A*.) *recki* Wainstein, *T*. (*A*.) *rhenanus* (Oudemans), *T*. (*A*.) *salvia* (Kolodochka), *T*. (*A*.) *spiralis* (Wainstein & Kolodochka), and mostly based on the shape of calyx of spermatheca and general apperance. Differences between *E. ueckermanni* **sp. nov.** and related species are given in Table 1.

**TABLE 1.**
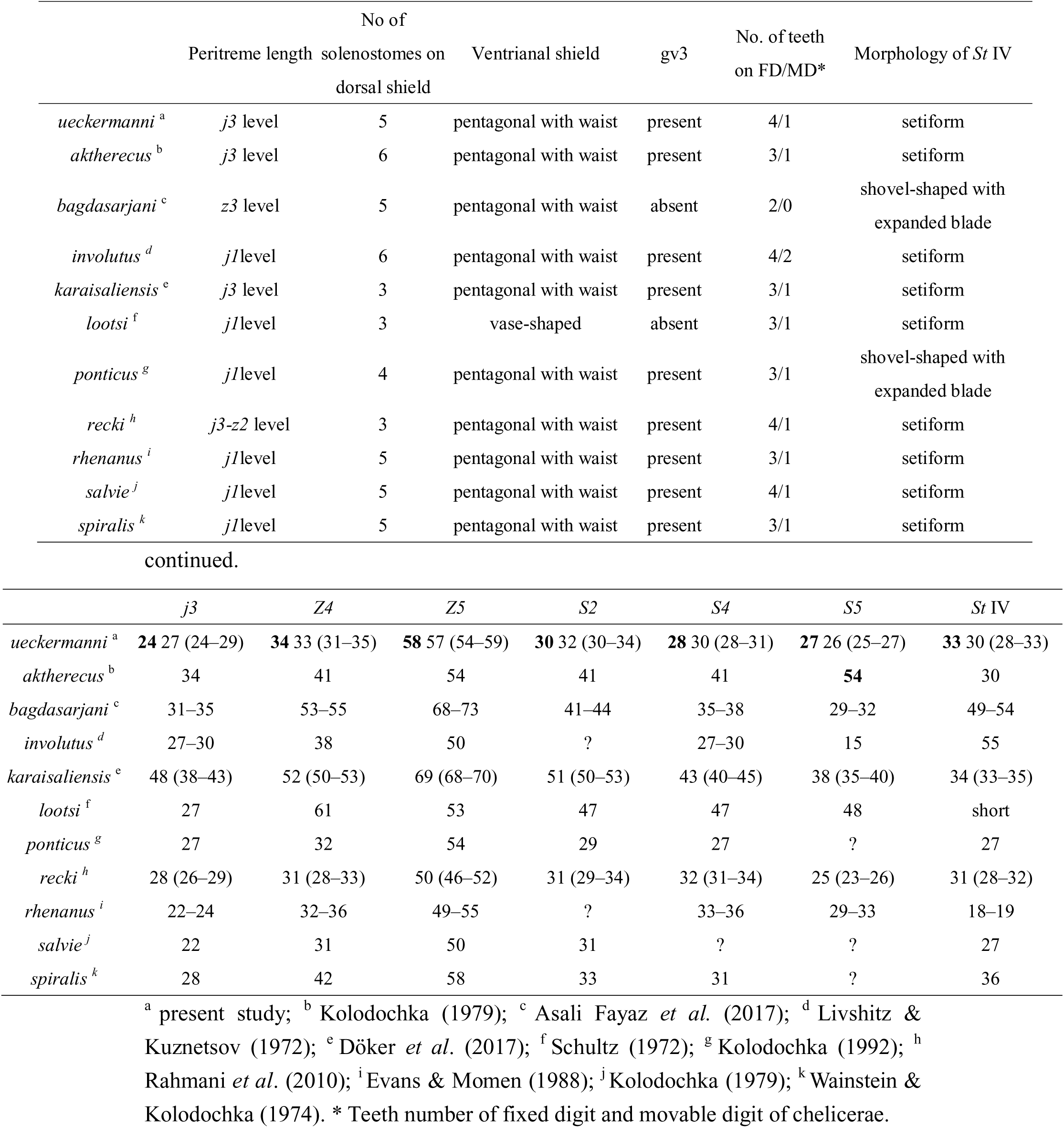
Differences between *Typhlodromus* (*Anthoseius*) ueckermanni sp. nov. and related species.

|  | Peritreme length | No of solenostomes on dorsal shield | Ventrianal shield | gv3 | No. of teeth on FD/MD* | Morphology of *St* IV |
| --- | --- | --- | --- | --- | --- | --- |
| *ueckermanni*<sup>a</sup> | *j3* level | 5 | pentagonal with waist | present | 4/1 | setiform |
| *aktherecus*<sup>b</sup> | *j3* level | 6 | pentagonal with waist | present | 3/1 | setiform |
| *bagdasarjani*<sup>c</sup> | *z3* level | 5 | pentagonal with waist | absent | 2/0 | shovel-shaped with expanded blade |
| *involutus*<sup>d</sup> | *j1* level | 6 | pentagonal with waist | present | 4/2 | setiform |
| *karaisaliensis*<sup>e</sup> | *j3* level | 3 | pentagonal with waist | present | 3/1 | setiform |
| *lootsi*<sup>f</sup> | *j1* level | 3 | vase-shaped | absent | 3/1 | setiform |
| *ponticus*<sup>g</sup> | *j1* level | 4 | pentagonal with waist | present | 3/1 | shovel-shaped with expanded blade |
| *recki*<sup>h</sup> | *j3-z2* level | 3 | pentagonal with waist | present | 4/1 | setiform |
| *rhenanus*<sup>i</sup> | *j1* level | 5 | pentagonal with waist | present | 3/1 | setiform |
| *salvie*<sup>j</sup> | *j1* level | 5 | pentagonal with waist | present | 4/1 | setiform |
| *spiralis*<sup>k</sup> | *j1* level | 5 | pentagonal with waist | present | 3/1 | setiform |

|  | *j3* | *Z4* | *Z5* | *S2* | *S4* | *S5* | *St* IV |
| --- | --- | --- | --- | --- | --- | --- | --- |
| *ueckermanni*<sup>a</sup> | **24** 27 (24–29) | **34** 33 (31–35) | **58** 57 (54–59) | **30** 32 (30–34) | **28** 30 (28–31) | **27** 26 (25–27) | **33** 30 (28–33) |
| *aktherecus*<sup>b</sup> | 34 | 41 | 54 | 41 | 41 | **54** | 30 |
| *bagdasarjani*<sup>c</sup> | 31–35 | 53–55 | 68–73 | 41–44 | 35–38 | 29–32 | 49–54 |
| *involutus*<sup>d</sup> | 27–30 | 38 | 50 | ? | 27–30 | 15 | 55 |
| *karaisaliensis*<sup>e</sup> | 48 (38–43) | 52 (50–53) | 69 (68–70) | 51 (50–53) | 43 (40–45) | 38 (35–40) | 34 (33–35) |
| *lootsi*<sup>f</sup> | 27 | 61 | 53 | 47 | 47 | 48 | short |
| *ponticus*<sup>g</sup> | 27 | 32 | 54 | 29 | 27 | ? | 27 |
| *recki*<sup>h</sup> | 28 (26–29) | 31 (28–33) | 50 (46–52) | 31 (29–34) | 32 (31–34) | 25 (23–26) | 31 (28–32) |
| *rhenanus*<sup>i</sup> | 22–24 | 32–36 | 49–55 | ? | 33–36 | 29–33 | 18–19 |
| *salvie*<sup>j</sup> | 22 | 31 | 50 | 31 | ? | ? | 27 |
| *spiralis*<sup>k</sup> | 28 | 42 | 58 | 33 | 31 | ? | 36 |

<sup>a</sup> present study; <sup>b</sup> Kolodochka (1979); <sup>c</sup> Asali Fayaz *et al.* (2017); <sup>d</sup> Livshitz & Kuznetsov (1972); <sup>e</sup> Döker *et al.* (2017); <sup>f</sup> Schultz (1972); <sup>g</sup> Kolodochka (1992); <sup>h</sup> Rahmani *et al.* (2010); <sup>i</sup> Evans & Momen (1988); <sup>j</sup> Kolodochka (1979); <sup>k</sup> Wainstein & Kolodochka (1974). \* Teeth number of fixed digit and movable digit of chelicerae.

### 21. Typhlodromus (Anthoseius) dossei Schicha, 1978

#### Material examined

Australia, one female (HAL095G077) from *Banksia* sp. (Proteaceae), 7 Dec 2006.

#### Distribution

Oceania: Australia (Schicha 1978)

### Remarks

The species has only been recorded in Australia (Demite *et al*. 2020). Schicha (1978) reported that the species fed on an eriophyid mite colony from *Ficus carica.* He also reported that this species was collected from various parts of Australia.

### 22. Typhlodromus (Anthoseius) recki Wainstein, 1958

#### Material examined

France, one female (QAR102H041) from *Thymes* sp. (Lamiaceae), 6 Sept 2013.

#### Distribution

Africa: Algeria (Athias-Henriot 1960), Tunisia (Kreiter *et al*. 2002). Asia: Armenia (Arutunjan 1970), Azerbaijan (Gadzhiev & Abbasova 1965), Cyprus (Papadoulis *et al*. 2009), Iran (Rahmani *et al*. 2010), Israel (Amitai & Swirski 1978), Kazakhstan (Wainstein 1958), Lebanon (Dosse 1967), Syria (Barbar 2016), Turkey (Swirski & Amitai 1982). Europe: Austria (Ragusa & Ragusa 1997), France (Kreiter & Brian 1987), Georgia (Wainstein 1958), Greece (Swirski & Ragusa 1977), Hungary (Bozai 1980), Italy (Castagnoli *et al*. 1984), Moldova (Kolodochka 1980), Portugal (Ferreira & Carmona 1994), Russia (Wainstein 1958), Slovenia (Kreiter *et al*. 2020), Ukraine (Livshitz & Kuznetsov 1972).

#### Remarks

The species is distributed in Africa and Europe (Demite *et al*. 2020), and only one specimen was collected from commodity of France. Papadoulis *et al*. (2009) provideda detailed redescription, and mentioned that the species prefers to inhabit Lamiaceae plants; we also found the species from a Lamiaceae plant.

### 23. Typhlodromus (Anthoseius) rhenanus (Oudemans, 1905)

#### Material examined

USA, one female (HAL095G065) from *Thymus vulgaris* (Lamiaceae), 2 Nov 2006; Netherlands, one female (HAL098B507) from *Rosmarinus officinalis* (Lamiaceae), 2 Oct 2009; Netherlands, one female (HAL100B067) from *Rosmarinus officinalis* (Lamiaceae), 19 May 2011; Netherlands, six females (HAL101B182) from *Rosmarinus officinalis* (Lamiaceae), 30 Mar 2012.

#### Distribution

Africa: Algeria (Athias-Henriot 1957b), Tunisia (Kreiter *et al*. 2002). Asia: Azerbaijan (Gadzhiev & Abbasova 1965), Cyprus (Georghiou 1959), India (Narayanan & Ghai 1961), Iran (Khalil-Manesh 1973), Israel (Swirski & Amitai 1961), Kazakhstan (Andreeva 1966), Syria (Barbar 2013), Turkey (Çobanoğlu 1989a). Europe: Belarus (Sidlyarevich 1966), Belgium (Nesbitt 1951), Denmark (Hansen & Johnsen 1986), England (Nesbitt 1951), Finland (Tuovinen & Rokx 1991), France (Nesbitt 1951), Germany (Oudemans 1905a), Greece (Papadoulis *et al*. 2009), Hungary (Kropczynska & Jenser 1968), Italy (Gunthart 1960), Latvia (Salmane & Petrova 2002), Moldova (Kolodochka 1980b), Montengro (Mijuskovic & Tomasevic 1975), Netherlands (Miedema 1987), Norway (Edland 1987), Poland (Suski 1961), Portugal (Carmona 1962), Russia (Beglyarov 1957), Serbia (Radivojevi & Petanović 1984), Slovakia (Fend’a 2010), Slovenia (Bohinc & Trdan 2013), Spain (Iraola *et al*. 1997), Sweden (Steeghs *et al*. 1993), Switzerland (Gunthart 1960), Ukraine (Kolodochka 1974). North America: Canada (Nesbitt 1951), USA (Nesbitt 1951). South America: Brazil (Diehl *et al*. 2012);

#### Remarks

The species is distributed worldwide, except in Oceania (Demite *et al*. 2020). Chant (1959) reported the species after Oudemans (1905) described it. Subsequent researchers provided conflicting opinions on the species status. Evans & Momen (1988) examined the type materials of the species and related species and provided a detailed redescription.

### 24. Typhlodromus (Anthoseius) vulgaris Ehara, 1959

#### Material examined

Japan, one female (HAL100B195) from *Vitis vinifera* (Vitaceae), 10 Nov 2011.

#### Distribution

Asia: China [Hong Kong (Swirski & Shechter 1961)], Iran (Khalil-Manesh 1973), Japan (Ehara 1959), South Korea (Ryu & Ehara 1990). Europe: Russia (Sapozhnikova 1966).

#### Remarks

The species is distributed in Asia and Europe (Demite *et al*. 2020). Kishimoto *et al*. (2013) reported that the species is considered as lifestyle Type III, generalist predator that could become an effective predator of spider mites and eriophyoid mites in field.

### 25. Typhlodromus (Typhlodromus) pyri Scheuten, 1857

#### Material examined

Netherlands, one female (HAL100B198) from *Rosmarinus officinalis* (Lamiaceae), 3 Nov 2011.

#### Distribution

Africa: Egypt (El-Badry 1967). Asia: Azerbaijan (Abbasova 1970), Saudi Arabia (Fouly & Al-Rehiayani 2011), Turkey (Çobanoğlu 1991). Europe: Austria (El-Borolossy 1989), Belarus (Sidlyarevich 1966), Belgium (Chant *et al*. 1974), Croatia (Tixier *et al*. 2010), Czech Republic (Hluchy *et al*. 1991), Denmark (Chant *et al*. 1974), England (Chant 1959), Finland (Tuovinen 1993), France (Rambier 1974), Germany (Scheuten 1857), Greece (Swirski & Ragusa 1977), Hungary (Kropczynska & Jenser 1968), Italy (Castagnoli & Liguori 1986), Moldova (Wainstein 1973), Montenegro (Mijuskovic & Tomasevic 1975), Netherlands (Van de Vrie 1963), Norway (Edland 1987), Poland (Wiackowski & Suski 1963), Portugal (Carmona 1962), Russia (Meshkov 1999), Serbia (Kropczynska & Petanović 1987), Slovakia (Jedlickova & Kolodochka 1994), Slovenia (Bohinc & Trdan 2013), Spain (Pérez Otero & Mansilla Vázquez 1997), Sweden (Chant *et al*. 1974), Switzerland (Ragusa & Swirski 1976), Ukraine (Kolodochka 1974). North America: Canada (Putman & Herne 1966), USA (Chant 1959). Oceania: Australia (Schicha 1987), New Zealand (Collyer 1964a). South America: Chile (Ragusa & Vargas 2002)

#### Remarks

The species is distributed worldwide (Demite *et al*. 2020), but only one specimens from the Netherlands has been found. EPPO (2020) listed this species as a commercially used biological control agent. The species is considered as lifestyle Type III-a, generalist predator that lives on pubescent leaves, and also has the ability to feed on fungi (McMurtry *et al*. 2013).

## Acknowledgements

We thank to BAPHIQ and inspectors who found the samples in plant quarantine. We also thank to E. Ueckermann (Pretoria, South Africa), İ. Döker (CU, Turkey), S. F. Lin (NCHU, Taiwan), Y. Hsiao (CSIRO & ANU, Australia), C. H. Chen (BAPHIQ, Taiwan), F. Y. Ning (BAPHIQ & NTU, Taiwan) for suggestions. Thank to H. Ono (NSMT, Japan), L.A. Corpuz-Raros and J. Naredo (UPLB-MNH) for borrowing type specimens for comparison. Thanks to Wallace Academic Editing for English editing of the draft. The study is supported by grants MOST 105-2621-B-002-002-MY3 and MOST 108-2621-B-002-005-MY3 from the Ministry of Science and Technology, Taiwan.

